# Drug combination prioritization for cancer treatment using single-cell RNA-seq based transfer learning

**DOI:** 10.1101/2022.04.06.487357

**Authors:** Daniel Osorio, Daniel J. McGrail, Nidhi Sahni, S. Stephen Yi

**Affiliations:** Department of Oncology, Livestrong Cancer Institutes, Dell Medical School, The University of Texas at Austin, Austin, TX 78712, USA; Center for Immunoterapy and Precision Immuno-Oncology, Cleveland Clinic, Cleveland, OH 44195, USA; Department of Epigenetics and Molecular Carcinogenesis, The University of Texas, MD Anderson Cancer Center, Houston, TX 77230 USA; Department of Bioinformatics and Computational Biology, The University of Texas, MD Anderson Cancer Center, Houston, TX 77030, USA; Quantitative and Computational Biosciences Program, Baylor College of Medicine, Houston, TX 77030, USA; Interdisciplinary Life Sciences Graduate Programs (ILSGP), College of Natural Sciences, The University of Texas at Austin, Austin, TX 78712, USA; Oden Institute for Computational Engineering and Sciences (ICES), The University of Texas at Austin, Austin, TX 78712, USA; Department of Biomedical Engineering, Cockrell School of Engineering, The University of Texas at Austin, Austin, TX 78712, USA

**Keywords:** synergistic-effect, drug-combination, precision-oncology, single-cell RNA-seq, transfer-learning

## Abstract

Precision oncology seeks to match patients to the optimal pharmacological regimen; yet, due to tumor heterogeneity, this is challenging. Numerous studies have been conducted to produce clinically relevant pharmacological response forecasts by integrating modern machine learning algorithms and several data types. Insufficient patient numbers and lack of knowledge of the molecular targets for each drug under study limit their use. As a proof of concept, we use single-cell RNA-seq based transfer learning to contextualize patients’ tumor cells in terms of their more similar cell lines with known susceptibility to drug combinations. Our objective is to maximize the translational potential of *in-vitro* assays for identifying synergistic drug combinations and prioritizing them for clinical use. Consistent findings in a cohort of breast cancer patients corroborated our understanding of the disease’s molecular subtypes. To aid in creating personalized treatments and data-driven clinical trials, we identified the most prevalent cell lines and prioritized synergistic combinations based on tumor compositions at various resolution levels.

## Introduction

Precision oncology strives to match patients with the best pharmaceutical regimen for treating their cancer (1). Due to the heterogeneity of malignancies, selecting the right pharmacological regimen is a difficult challenge (2). Tumors are made up of a variety of cell types that interact and modify the extracellular matrix, influencing the immune system’s ability to recognize and kill malignant cells (3). For that reason, cancer patients are usually categorized into subgroups based on histological or molecular characteristics such as cell morphologies, DNA mutations, or gene expression patterns to improve treatment efficacy (4, 5). Using this approach, only patients having characteristics that are comparable to those of others who have had a positive response to treatment are given the medicine (6).

Several research groups have recently attempted to incorporate multiple sources of information into cutting-edge machine learning approaches in order to create clinically oriented predictions of patient responses to medications. I-PREDICT (7), NCI-MATCH (8), MI-ONCOSEQ (9), WINTHER (10), ONCO-TARGET/ONCOTREAT (11), SELECT (12), and ENLIGHT (13) are studies that aim to match patients with therapies and predict their impact on treatment outcome using DNA biomarkers, genomic and transcriptomic information, protein-protein interaction networks, synthetic lethal and/or synthetic rescue interactions among other factors (7–13). However, developing such studies requires data from a large number of patients as well as extensive knowledge of the exact molecular targets for each of the drugs under investigation, limiting their applicability (14).

*In-vitro* cell line assays, on the other hand, are still the first line of cancer research since they serve as surrogates for the patient’s tumor response. (15). Cancer cell lines are a low-cost, easy-to-maintain model system for medical research suitable to be manipulated to represent the genetic aberrations found in patients’ tumors (16, 17). When integrated into panels, cell lines represent the genetic diversity and phenotypic variability of the patients; hence, it is necessary to develop strategies to optimize the translational potential of *in-vitro* cellular responses into pharmaceutical alternatives for patients without the need of a reference group (18).

With the introduction of new high-throughput experimental approaches and computational methods that allow for the characterization of the transcriptional profile at the single-cell level, we can now identify isogenic sub-populations of cells responding differently to the drug under study (19). When applied to tumors, single cell approaches allow identifying of all the proportions of cell types making part of it (20), and when both sources of information (responses of purified cell lines and tumor data) are combined, single-cell transcriptomics offers the ability to maximize the translational potential of *in-vitro* cellular responses into patients’ pharmacological options.

Recently, Gambardella and collaborators released a single-cell atlas of breast cancer cell lines (21). The atlas reports single-cell RNA-seq data for 35, 276 individual cells from 32 breast cancer cell lines individualized using DROP-seq (22). Concurrently, using 10× Chromium (23), Wu and collaborators published a single-cell atlas of human breast cancers that includes 130, 246 annotated single cells from 26 primary tumors, including 11 ER^+^, 5 HER2^+^ and 10 TNBCs (24). Together, both data sets provide an opportunity to contextualize the cancer cells from patients’ tumors in terms of cancer cell lines using transfer learning. Transfer learning is a machine learning technique that uses a joint embedding to map query data sets on top of a given reference (25). This method allows to integrate datasets after removing the non-biological variation and to contextualize n data sets using the metadata associated in the existing reference with high precision and without the need of recomputing the reference embedding (26, 27).

The ability to contextualize tumor cells in terms of more similar cell lines opens up a world of possibilities for mining hundreds of publicly available cell line assays to identify appropriate pharmaceutical options for treating patients in a tailored manner without using a reference cohort or having a deep understanding of the drugs’ molecular targets. Additionally, this strategy may be broadly used to any illness for which a sufficient number of cell lines representing patients’ genetic and phenotypic heterogeneity as well as *in-vitro* assays are available.

Here, as a proof of concept, we used the systematic evaluation of 1, 275 pairwise drug combinations tested over 51 breast cancer cell lines using the Genomics of Drug Sensitivity in Cancer (GDSC) cell line screening platform to prioritize synergistic drug combinations based on patients’ tumor composition aiming to improve patients’ survival rate through personalized medicine (28). We also examined the data at the population level to provide reference of which cell lines and drug combinations prioritize for drug-development and clinical trial purposes in each breast cancer molecular subtype. We were able to trace tumor cell transcriptional profiles to their most similar derived from cell lines using the transfer learning data-driven computational approach as a compass. We provided evidence that transfer learning is valuable for precision medicine because it bridges the gap between *in-vitro* drug sensitivities and pharmacological treatment options for a given patient or patient subgroup.

## Methods

### Single-cell RNA-seq datasets

We compiled a collection of publicly accessible single-cell RNA-seq count matrices from a variety of sources (See Data Availability). When needed, gene identifiers (IDs) were translated into the current gene symbols using the reference provided by ENSEMBL BioMart for GRCh38.p13 (29). The data were not subjected to any further normalization, transformation or quality control filters.

### Drug sensitivity data

We downloaded the viability data of 51 breast, 45 colorectal, and 29 pancreatic cancer cell lines in response to 2, 025 clinically relevant drug combinations reported through the the Genomics of Drug Sensitivity in Cancer (GDSC) cell line screening platform (28). We filtered the drug combinations evaluated in breast cancer cell lines and used this information to prioritize combinations at various resolution levels according to tumor composition.

### Reference atlas construction and validation

The low-dimensional embedding representing the transcriptional profile of 32 breast cancer cell lines cultured independently in ATCC recommended complete media at 37°C and 5% CO_2_ was constructed using Symphony (26). Symphony creates efficient and precise single-cell reference atlas through reference compression that extracts and organizes data from reference datasets into a unified and simple format that may be used to map query cells. In brief, cells are allocated in soft-cluster memberships repeatedly using Harmony, which computes weights for a linear mixture model in order to eliminate covariate-dependent effects (30). Symphony then compresses the reference into a mappable object, utilizing the reference-learned model parameters allowing to augment the embedding with additional query cells. It maps cells into the reference without recomputing, maintaining the structure of the reference atlas. Given that all of the reference cell lines employed in this study originated from the same project (21). The atlas was constructed using a constant covariate, which enabled Symphony to recognize transcriptional differences across cell lines and cluster the cells appropriately. The top 50 hamonized dimensions were chosen for reference compression, and all other parameters were left unchanged.

We performed cross-validation, selecting a varied number of cells (from 1, 000 to the size of the dataset with increasing steps of 1, 000 cells) to recompute the reference and map the cells with known labels that were left out to the reference. The performance was assessed using the (accuracy) percentage of correctly classified observations divided by the total number of observations through the ‘caret’ R package (31). We also used single-cell RNA-seq data from other laboratories and generated using a different single-cell technique to evaluate the performance of mapping three distinct breast cancer cell lines in their wild-type form, with a genetic change, and under the influence of an anti-cancer drug (32–34).

### Transfer learning approach and validation

We obtained the transcriptional profile of 130, 246 annotated single cells from from 26 primary breast cancer tumors (24). The authors of the dataset identified cells as malignant based on copy number variations (CNVs), and these cells were utilized to map them into the reference embedding of breast cancer cell lines. To determine the most equivalent cell line, we performed a *k*-nearest neighbors search of the closest five cells following cell mapping and assigned the cell to the label with the highest relative frequency. To evaluate the performance of the assignation, we used leave-one-out cross validation and reported the associated error for the calculated patient’s tumor composition in terms of breast cancer cell lines.

### Activity score assignation based on patients’ tumor composition

Equation 1 was used to rank synergistic drug combinations based on the patient’s tumor composition in terms of breast cancer cell lines. The activity score (*A*) of a two-drugs (*i* and *j*) combination is computed as the scaled (between −1 and 1) average of the synergistic effect (*S*) of both drugs across different concentrations (*d*) multiplied by the proportion (*P*) of the cell line (*l*) in tumor’s composition

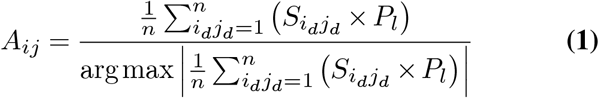

### P-value assignation based on cell lines’ response stability and patients’ tumor composition

To account for the uncertainty of the cell line response to the drug combination, the authors of the dataset reported the root-mean-square error (RMSE) of the measurement. We used this value to compute an unscaled activity score (*A*) penalized by the RMSE of the synergistic response to the combination 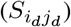 at different concentrations (*d*) using Equation 2.

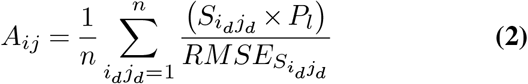

This penalized score was then used to produce a *P*-value based on the empirical distribution after 1, 000 bootstrap iterations.

### Drug combination prioritization

Prioritization of drug combinations was based on the computed activity score and associated P-value accounting for the tumor composition in terms of cell lines at various degrees of resolution, ranging from a single patient to the whole cohort of patients with breast cancer. The top drug combinations identified in each scenario were labeled. The Wilcoxon rank sum test was used to compare activity scores across breast cancer molecular subgroups.

## Results

### Overview of the drug combination prioritization procedure

Our strategy for optimizing the translational potential of *in-vitro* tests for discovering synergistic drug combinations and selecting them for clinical application is divided into four general steps as described in the Figure 1.

**Fig. 1.**
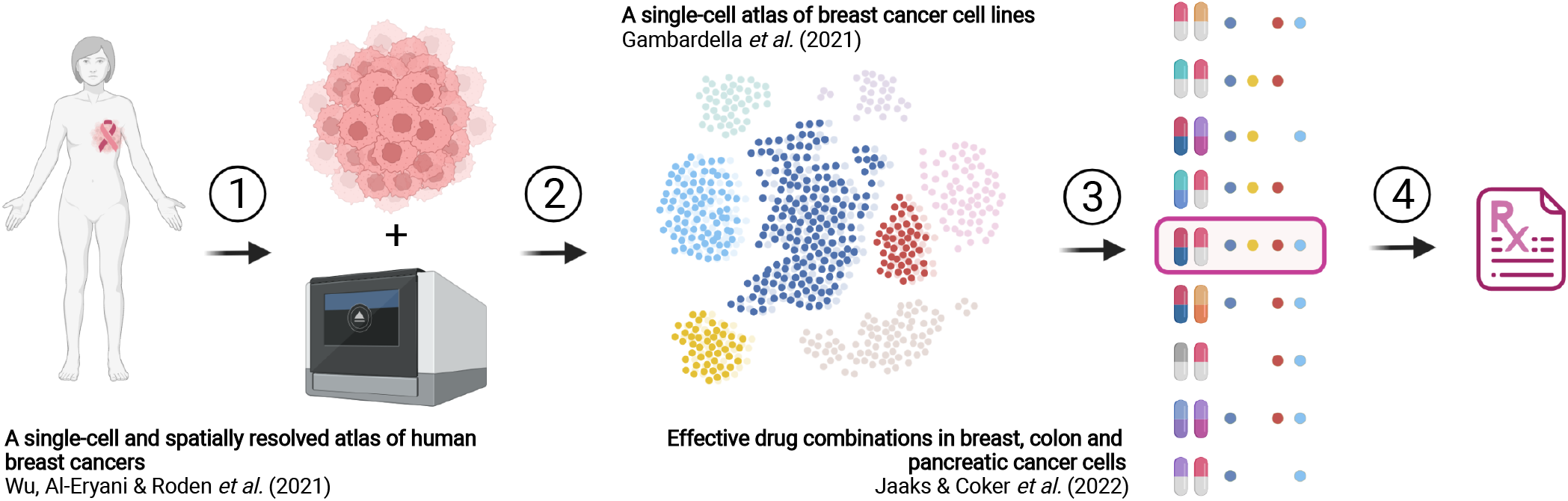
Approach overview. **(1)** A biopsy from the patient tumor is collected and single-cell transcriptomes are sequenced using drop-based single-cell RNA-seq techniques. **(2)** Using transfer learning, cells are mapped to an atlas of cancer cell lines derived from the tumor of origin and representing the phenotypic diversity of the malignancies affecting patients. **(3)** Based on the detected tumor composition in terms of cell lines, the drug combinations with highest synergistic activity across the detected cell lines is identified and then **(4)** suggested for its prescription to the patient.

The first step is to sequence the transcriptome of cancerous tumor cells obtained from patients with the disease; in this study, we focused on breast cancer patients due to the availability of public data. The second step is to map the patient’s cells to a more closely related cell line using a cell line atlas designed to represent the disease and contextualize tumor cells in terms of cell lines. In the third step, we leverage the transfer learning process from step 2 to close the gap between *in-vitro* drug sensitivities and pharmacological treatment options for the patient. This step assigns a ranking to the drug combinations based on their effect on the cell lines identified previously. Finally, drug combinations that are expected to be effective in treating the patient are recommended for their prescription.

### Generation of a reference breast cancer cell atlas embeeding

We downloaded 35, 276 transcriptomes from 32 ATCC-authenticated breast cancer cell lines that had been individualized using DROP-seq and used them to generate a reference embedding as described in the methods section (22). After compressing the transcriptional data, Symphony generated a uniform manifold approximation and projection (UMAP) using the top 50 PCA components that discriminates the 32 clusters of cells in the low-dimensional space with a minimal overlap (*<* 1%, Figure 2A), which is better to differenciate cell lines than the projection reported by the dataset’s original authors (21, 26). This indicates that the logarithm of the counts per million *log*(CPM + 1) normalization provides a greater ability to distinguish between breast cancer cell lines than the gene frequency–inverse cell frequency (GF-ICF) normalization method (35).

**Fig. 2.**
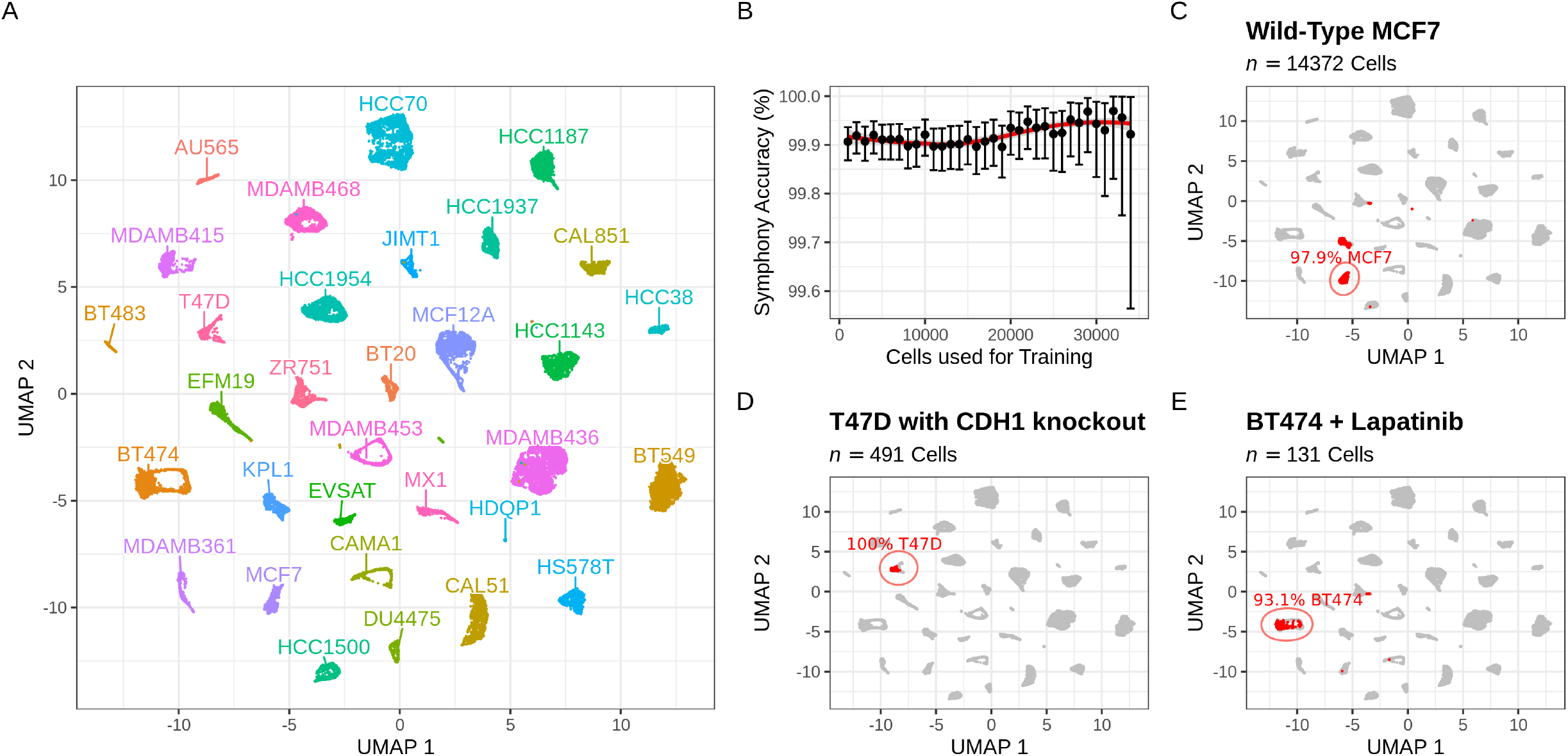
Transfer-learning accuracy evaluation. **(A)** Uniform Manifold Approximation and Projection (UMAP) displaying the harmonized transcriptional profile of 32 breast cancer cell lines sequenced using DROP-seq. **(B)** Accuracy evaluation using cross-validation. The percentage of cells utilized for training is displayed, as well as the accuracy and 95% confidence intervals computed using the binomial test. Accuracy is defined as the percentage of correctly classified observations divided by the total number of observations. **(C)** Accuracy evaluation using wild-type cells from a different single-cell technology. Transcriptomes of cells derived from the wild-type MCF7 cell line that were sequenced using 10× Chromium were appropriately labeled in 97.9% of cases. **(D)** Accuracy evaluation using cells from a different single-cell technology carrying a genetic modification. Transcriptomes of cells originating from the T47D cell line carrying the *CDH1* gene knockout through CRISPR and sequenced using 10x Chromium were labeled correctly in all cases. **(E)** Accuracy evaluation using cells from a different single-cell technology under cancer treatment. Transcriptomes of cells obtained from the BT474 cell line that were treated with 1 *µ*M of Lapatinib for 10 days and sequenced using 10x Chromium were identified correctly in 93.1% of instances.

### Evaluation of the transfer learning accuracy

We used two approaches to test the ability to distinguish between cell line transcriptomes. First, we used cross-validation to reconstruct the reference embedding of breast cancer cell lines using a randomly selected subset of cells (starting with 1, 000 and increasing by 1, 000 cells until we reached the entire dataset) and then remapped the portion of the cells with known labels that were left out to the reference embedding. Second, we mapped external transcriptomes from three different cell lines included in the breast cancer cell atlas that had been sequenced by other laboratories using a different single-cell RNA-seq technique to the reference embedding generated using all 35, 276 cells. In both cases, after remapping, the label for the cells were assigned using a *k*-nearest neighbors search of the closest five cells and selecting the label with the highest relative frequency.

The cross-validation approach’s results support the use of this technique in clinical settings. In all cases, we recovered the correct cell line label with greater than 99.5 percent accuracy (Binomial test, *P <* 0.01 in all cases, Figure 2B). However, these results may be distorted due to the fact that all transcriptomes used to generate the reference embedding and to assess performance were sequenced in the same batch. To avoid bias, we evaluated the performance of mapping transcriptomes from three distinct breast cancer cell lines included in the reference embedding in their wild-type state, with a genetic mutation, and under the influence of an anticancer drug sequenced using a different single-cell RNA-seq technique and by other laboratories (32–34).

To begin, we mapped 14, 372 single-cell transcriptomes sequenced using the 10× Chromium method from the wild-type MCF7 cell line into the reference embeeding of breast cancer cell lines to assess the efficacy of mapping transcriptomes from a different single-cell RNA-seq approach (32). We found that 14, 073, (97.92%) of them were recognized and labeled correctly as MCF7 (Figure 2C). The remaining cells were predominantly labeled as KPL1 (278 cells, 1.93%), and ZR751 (16 cells, 0.11%), two other estrogen receptor positive luminal cell lines linked to the same molecular subtype of breast cancer as MCF7 (36, 37). Furthermore, contamination of the KPL1 cell line by an MCF7 derivative has been documented (38); if this is the case for the parental cell line used by the dataset’s authors, then the cell label assignment may be considered as 99.85% accurate.

Because cancer cells have a high mutational load (39). We investigated the impact of carrying a genetic mutation related with epithelial cell phenotypic identity throughout the mapping process. We mapped 491 cell transcriptomes obtained from the T47D cell line bearing *CDH1* deletion created using CRISPR techniques for this purpose (33). We found that all 491 (100%) cells correctly assigned to the T47D cluster in the reference embeeding (Figure 2D), showing that genetic alterations, even those linked with phenotypic identity in breast cancer cells, seems to have little effect on the accuracy of our classifier.

Additionally, changes in anticancer drug response constitute a significant source of variability in cancer patients’ cell transcriptomes (40). As a result, we mapped 131 transcriptomes obtained from the BT474 cell line after ten days of treatment with 1*µ*M of Lapatinib (34). This was done to determine the influence of anticancer drugs on the cell identification during the mapping procedure. We found that 122 (93.13%) of the cells accurately mapped to the BT474 cluster (Figure 2E) in the reference embeeding. The remaining cells were attributed to ZR751 (8 cells, 6.11%) and CAMA1 (1 cell, 0.76%). Two metastatic estrogen receptor-positive luminal cells that are not of the same molecular subtype (Her2^+^) as BT474 (36, 41, 42). This finding provide evidence supporting that anticancer drugs affect the identity of cellular transcriptomes (43), and slightly reduce our ability to map treated cells’ transcriptomes into the reference embedding of breast cancer cell lines.

In general, when applied to transcriptomes derived from the same cell lines, the mapping approach and transfer learning procedure provide reliable results with accuracies above 90% in all cases, even when they were characterized using a different single-cell RNA-seq technique, affected by a genetic mutation, or undergoing treatment with an anticancer drug. All of these evidence supports its usage in clinical practice for classifying patients’ malignant tumor cells based on their most comparable cell line.

### Application of transfer learning to patients’ tumor cells

We obtained the transcriptomes of 30, 246 single cells from 11 ER^+^, 5 HER2^+^, and 10 TNBC primary tumors and selected the cells designated as malignant by the dataset’s authors using CNV characterization (24). After filtering the non cancerous cells, we kept the transcriptomes for 24, 489 cells from 20 tumors (9 ER^+^, 3 Her2^+^, and 8 TNBC). We used the transcriptomes of each patient’s associated malignant cells to map them into the reference embedding atlas established using breast cancer cell lines. This was done in order to contextualize the patient’s cells in form of their more similar cell lines. Here, we use the 894 cancer cells reported for donor CID44971 as an example of the process. When we mapped the cells transcriptomes into the reference embeeding, we found that 420 cells (46.98%) were mapped to MX1, 283 (31.66%) to HCC1187, 105 (11.74%) to CAMA1, 63 (7.05%) to ZR751, 15 (1.68%) to T47D, 2 (0.22%) to BT474, CAL51, and HS578T, and 1 (0.11%) to MDAMB436 and MDAMB468 (Figure 3A). This specific distribution across cell lines is unlikely to happen by chance (*χ*^2^ test, *P <* 1 *×* 10^*−*6^).

**Fig. 3.**
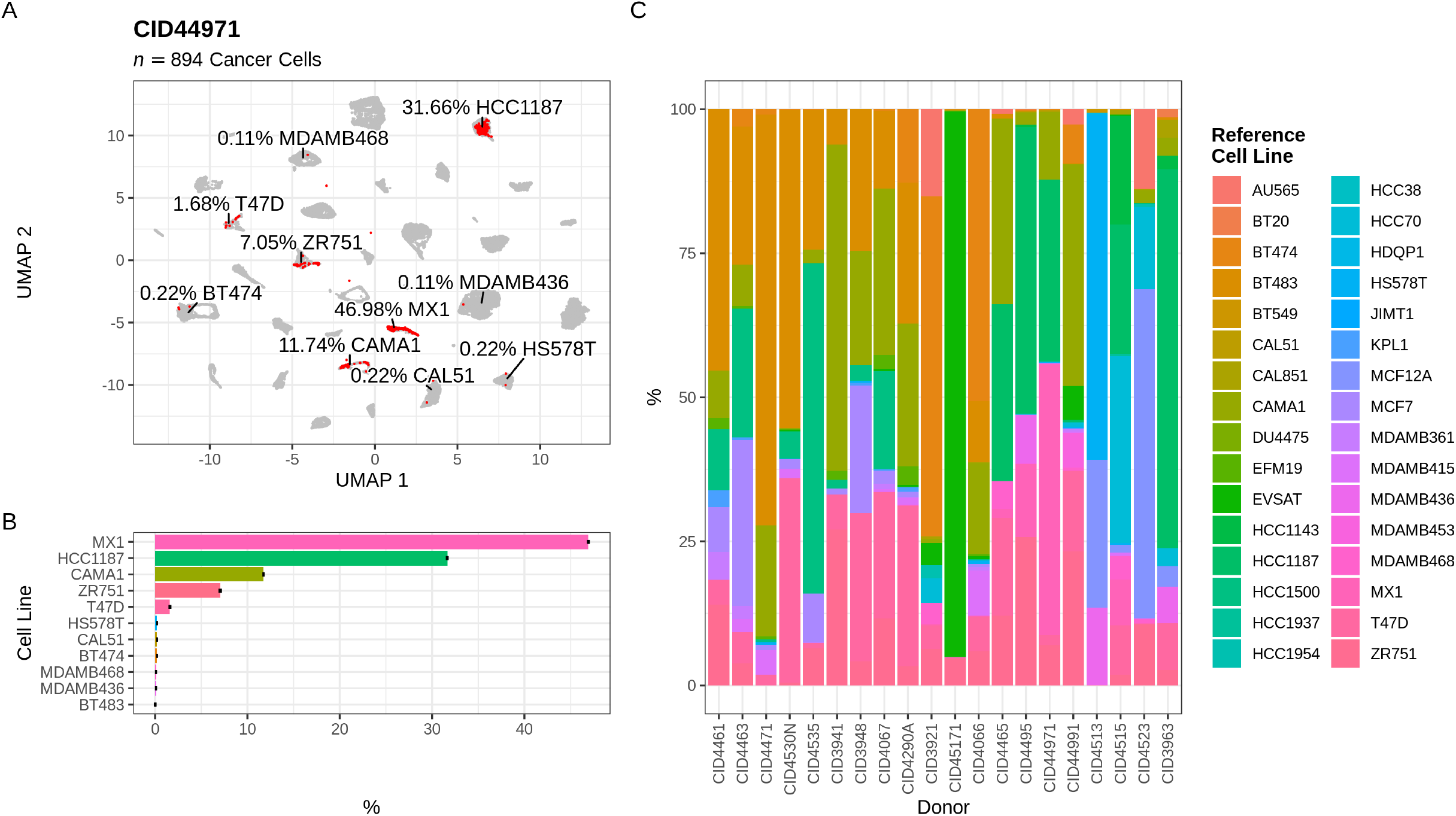
Determination of patient’s tumor cell composition in terms of cell lines. **(A)** Transcriptional profiles of the patient’s tumor cells mapped to their more comparable transcriptional profile within the integrated reference population of breast cancer cell lines. **(B)** Validation of the accuracy of the mapping procedure by leave-one-out cross validation (LOOCV). The CID44971 donor’s cancer cells were used, and the average and standard error of the relative frequency for each cell line are reported in percentages. **(C)** Composition of cancer tumor cells in terms of cell lines for twenty of the patients described in the breast cancer cell atlas.

Metadata associated to the dataset revealed that CID44971 cells are from a female donor affected with a breast cancer tumor of the TNBC molecular subtype. Confirming that diagnosis and the accuracy of the mapping and transfer learning approach, 709 (79.31%) of the cells’ transcriptomes were mapped to cell lines representing the TNBC molecular subtype, the remaining 185 cells were identified as derivates of luminal (183 cells, 20.47%) and Her2^+^ (2 cells, 0.22%) cell lines. A leave-one-out cross validation analysis was performed to validate the stability of the cell line assignation for the transcriptomes derived from the CID44971 donor (Figure 3B). The cross validated mean proportions were found to be highly correlated with those identified initially (*ρ* = 0.97, Spearman’s rank correlation *P* = 5.5 × 10^*−*20^) with small RMSE (0.02).

The computed cancerous cell proportions in terms of cell lines for all 20 donors are displayed in Figure 3C and numerically in Supplementary Figure S1. As previously described, cancer tumor cells are highly heterogeneous, facilitating the development of resistance to cancer therapies (44). The observed interpatient and intratumor heterogeneity, as seen in this patient cohort, makes drug development and clinical trial design difficult (45). Identifying the most common cell lines found in patients’ tumors across different molecular subtypes may aid in resource allocation and the development of treatments that will benefit a large proportion of the patient population (46).

### Prioritizing cell lines for breast-cancer drug development and testing

We averaged the relative frequencies identified after computing the cancerous cell proportions in terms of cell lines for all 20 donors to identify the most common cell lines found in patients’ tumors by molecular subtype. We found that the cell lines in which the cells from patients’ tumors were mapped are enriched with those developed as surrogates for each molecular subtype, confirming again the accuracy of the mapping and transfer learning approach. Luminal cell lines were shown to be abundant in estrogen receptor positive tumors, accounting for 95.79% of malignant cells (*χ*^2^ test, *P* = 8.46 ×10^*−*87^). We also observed that 77.46% of cells in Her2^+^ tumors mapped to cell lines designed to represent this category (*χ*^2^ test, *P* = 1.02 ×10^*−*130^), but only 56.22% of cells in TNBC mapped to cell lines representing this molecular subtype of breast cancer (*χ*^2^ test, *P* = 3.29 × 10^*−*44^). TNBC tumors were also found to be infiltrated with a modest proportion of luminal cell lines (28.07%) and a small proportion of Her2^+^ cell lines (4.72%).

Current breast cancer research relies on a small number of cell lines, with MCF7, T47D, and MDAMB231 accounting for more than two-thirds of all cell lines used in studies, raising the question of how representative these few cell lines are of the diversity of breast tumors with distinct clinical characteristics (47). This level of resolution, which enables us to classify patients’ tumor cells into cell lines, is relatively recent and the result of enormous advances in both experimental and computational biology (48). Here, we aim to characterize the predicted subpopulations of cell lines found in patients’ tumors. In contrast to conventional wisdom, we found that BT483 and CAMA1 are the most frequently occurring cells in estrogen receptor positive tumors, summing more than 50% of cells. For Her2^+^ tumors, we identified that BT474 and EVSAT are the two most prevalent, accounting for roughly 70% of tumor cells. The high frequency of BT474 in Her2^+^ tumors has previously been reported (33), lending additional support to the mapping and transfer learning approach’s high accuracy. In the case of TNBC tumors, we did not find a pair of cell lines that accounted for the vast majority of the tumor’s cells. HCC1187 and MX1 are the two most common cell lines, accounting only for 33.8 percent of tumor cells (Figure 4).

**Fig. 4.**
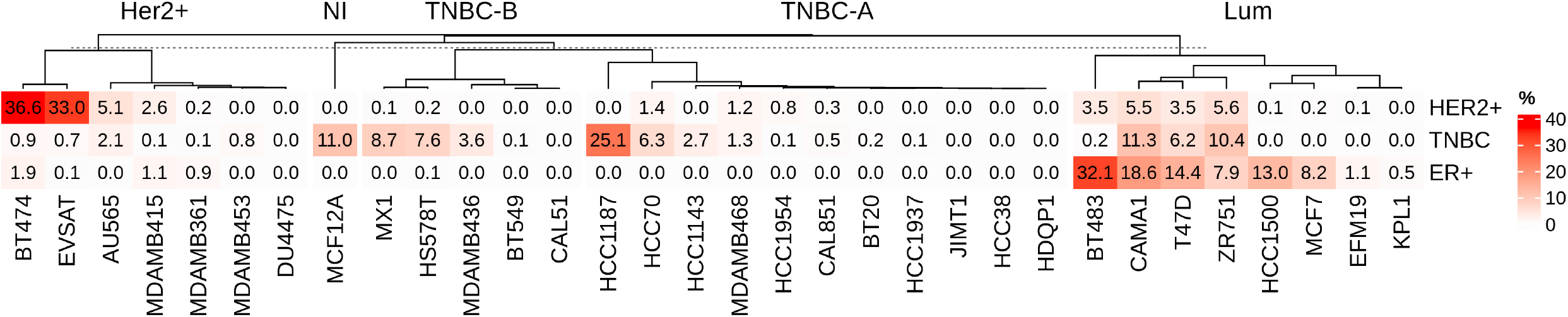
Identification of the tumor compositions of a population of patients. Composition in percentages of cancer tumor cells in terms of cell lines for twenty patients reported in the breast cancer cell atlas is summarized by molecular subtypes.

Cancer is a significant public health problem; it is currently the second leading cause of death for men and women in the US, passed only by heart disease (49). Aware of the limitations associated with the adoption of single-cell RNA-seq characterization of patients in clinical practice, we assessed the performance of various approaches for recapitulation of the high-resolution single-cell RNA-seq data, with the goal of leveraging molecular profiles from large cohorts already available, such as the TCGA project (50). Deconvolution of bulk RNA-seq profiles from a large cohort of patients into cell lines will provide unbiased knowledge for resource prioritization in cancer research with a high impact on public health. Deconvolution methods have demonstrated promising results in classifying heterogeneous tissues into distinct cell types (51, 52). However, their utility in identifying sample fractions using homogeneous cell types with minor molecular phenotype changes as reference has not been established.

We constructed a pseudo bulk sample by randomly selecting 1000 transcriptomes from the pool transcriptomes available for the 32 breast cancer cell lines. As a reference, we used the sum of all UMIs for all cells in each cell line. Following that, both the constructed sample and the reference were normalized using the logarithm of the counts per million. We used CIBERSORTx (51) to deconvolve the constructed sample and recover the fraction of cell lines whose true proportion is known. The result is a negative correlation (*ρ* = − 0.45, Spearman’s rank correlation, *P* = 9.33 × 10^*−*3^) between the predicted and known cell composition (Supplementary Figure S2). We also tried using Bisque (52), but this tool requires paired bulk and single cell RNA-seq data for a portion of the population in order to generate a reference expression profile and learn gene-specific bulk expression transformations for robust RNA-seq data decomposition. Due to the lack of such pairing profiles, their application to the TCGA data was not feasible.

Driven by the fact that deconvolution did not produce useful results. We used Spearman’s correlation to determine whether clustering enabled us to determine at least the most frequent cell line making part of the tumor samples (53). To accomplish this, we computed the pseudobulk profile, which includes all cells in the tumor (for reference in cases where FACS is not possible) and only cancerous tumor cells. The generated pseudobulk profiles were normalized using the logarithm of the counts per million and then combined with the cell line pseudobulk profiles. The normalized pseudo bulk profiles of samples and cell lines were clustered, and the association’s uncertainty was evaluated using bootstrap through ‘pvclust’ (54). In both cases, whether we consider all cells within the tumor sample (Supplementary Figure S3) or only cancerous cells (Supplementary Figure S4), the Spearman’s correlation fails to recognize the most frequently occurring cell line within the tumor samples.

The ability to contextualize patients’ tumor cells in terms of more similar cell lines opens up a world of possibilities for mining hundreds of publicly available cell line assays to identify appropriate pharmaceutical options for treating patients in a personalized manner without the need for a reference cohort or a thorough understanding of the drugs’ molecular targets. To determine the best pharmacological regimen for each patient based on the cell line composition of their tumor, a collection of phenotypic responses (for example, survival) to drugs across cell lines is all that is required. However, because monotherapy in cancer is highly susceptible to resistance development after an initial response to treatment (55). Combination therapy has become the standard pharmacological regimen for treating complex diseases like cancer. Combination therapy improves patient survival by halting tumor progression and preventing the development of drug resistance in cancer (56). As a result, we focused on identifying drug combinations with synergistic effects and ranking them based on the tumor composition of the patients.

### Prioritizing synergistic drug combinations for personalized medicine and clinical trials

We downloaded the 296, 707 survival statistics generated from the testing of 2, 025 clinically relevant two-drug combinations in 51 breast, 45 colorectal, and 29 pancreatic cancer cell lines (28). After filtering those from breast cancer cell lines, we ended with 156, 065 records repre-senting the effect of 1, 275 two-drug combinations at different concentrations. To evaluate the proposed activity score’s performance, we computed it using a uniform proportion for all cell lines representing each breast cancer subtype (Supplementary Figure S5). We found that the combination of Cisplatin and Gemcitabine is recommended as the best candidate for TNBC patients under this setting, confirming its accuracy. Cisplatin and Gemcitabine are used as first-line therapy in patients affected with metastatic triple negative breast cancer (57).

Once its performance was confirmed, the activity score was computed for each patient and molecular subtype of breast cancer, as well as for the entire cohort, as described in Equation 1, using the identified proportions for each cell type in each case. As expected given the diversity of tumor compositions, the top candidates for each molecular subtype and patient were different (Figure 5). In consistency with the reported by the authors of the dataset, drug combinations including Navitoclax, – an experimental orally active *BCL2, BCL2L1*, and *BCL2L2* inhibitor (58)–are enriched among the top candidates displaying synergistic effects for tumors from the entire cohort of patients (Hypergeometric test, *P* = 5.30 × 10^*−*5^) and for the estrogen receptor positive (Hypergeometric test, *P* = 9.53 × 10^−7^) and Her2^+^ (Hypergeometric test, *P* = 5.30 × 10^−5^) molecular sub-types of breast cancer (Figure 5, Panels B–D), but not for TNBC tumors (Figure 5E). This result is directly related to the unique tumor compositions found, rather than a generalized response of TNBC cell lines to Navitoclax combinations (Supplementary Figure S6).

**Fig. 5.**
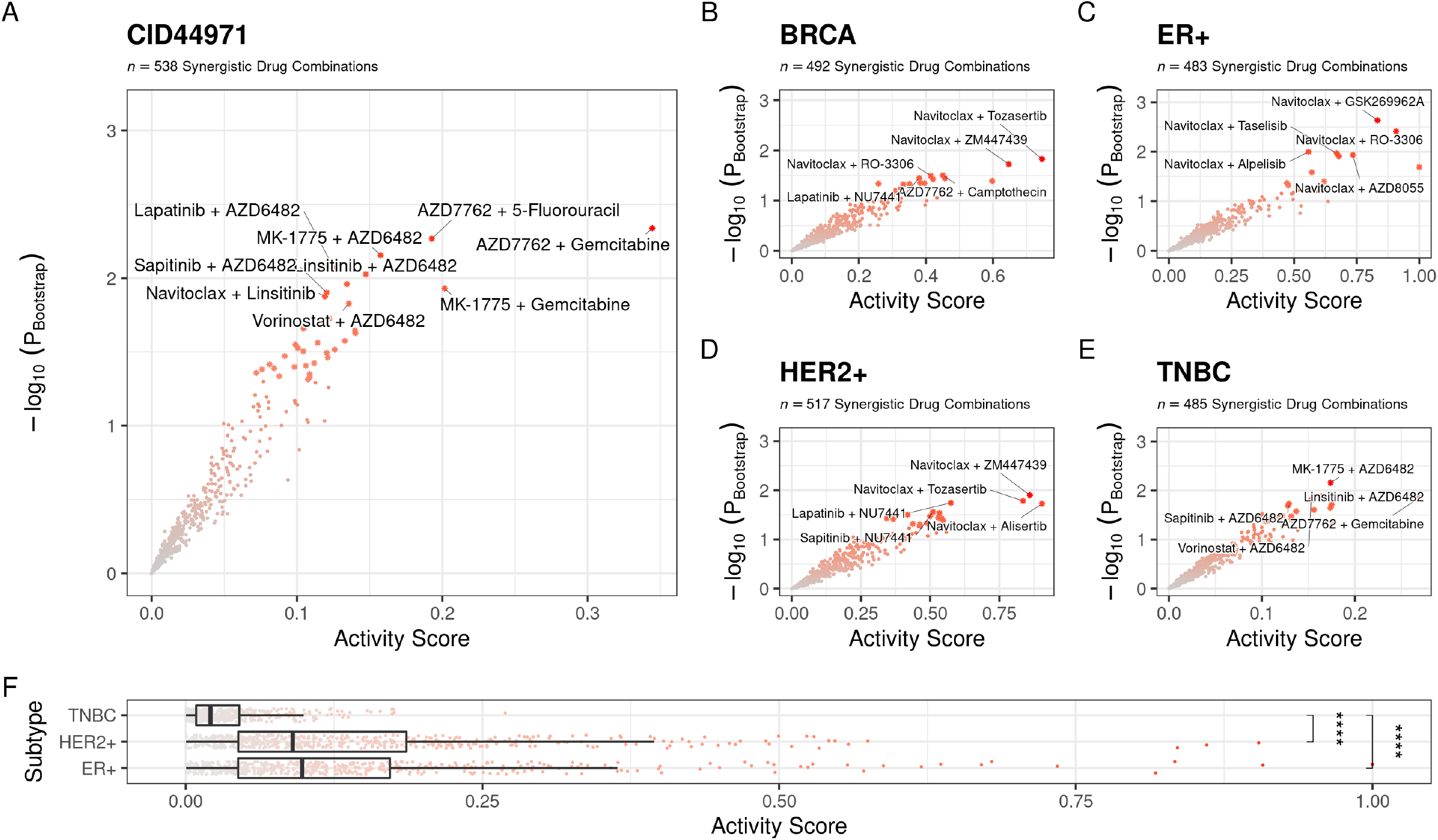
Prioritization of drug combinations exhibiting synergistic effects based on the tumor composition in terms of cell lines. **(A)** Computed for the CID44971 donor. **(B)** Computed for all donors included in the breast cancer atlas. **(C)** Computed for donors with ER-positive malignancies. **(D)** Computed for donors with HER2-positive malignancies. **(E)** Computed for donors with TNBC malignancies. **(F)** Comparison of the computed activity score across breast cancer sub-types.

We found that the best candidate for TNBC malignancies was the combination of AZD7762 and Gemcitabine, which has been shown to be effective in triple negative breast cancer (59). However, development of AZD7762 has been suspended due to significant heart side effects (60). We also found an enrichment of drug combinations having AZD6482 making part of it (Hypergeometric test, *P* = 1.9 ×10^−9^). AZD6482 is an ATP competitive inhibitor of the phosphatidylinositol 3-kinase (*PI3K*) p110*β* isoform with an IC_50_ of 0.01 nM (61). *PI3K* is an important target for breast cancer, because *PTEN* loss (the sixth most common mutation in breast cancer patients’ tumors) causes upregulation of *PI3K*/*AKT*, through the p110*β* isoform (62, 63). When AZD6482 is combined with Linsitinib (selective inhibitor of *IGF1R*), Sapitinib (a reversibles ATP competitive inhibitor of *EGFR, ERBB2* and *ERBB3*), Vorinostat (an HDAC inhibitor) or with MK-1775 (also known as Adavosertib, a selective *WEE1* inhibitor), it display singergistic effects against breast cancer cell lines (28). These findings support AZD6482’s versatility for the treatment of triple negative breast cancer and provide options for managing resistance development via four distinct pathways.

When we compared all of the computed activity scores across molecular subtypes of breast cancer, we discovered that those computed for TNBC are significantly lower (Figure 5F) than those computed for HER2^+^ (ANOVA, *P* = 2.27 ×10^−4^) and ER^+^ subtypes (ANOVA, *P* = 1 ×10^−7^). Finding effective treatments for TNBC has been a difficult task; here, we provide evidence that such difficulty is highly associated with the hyper-heterogeneity of the cancerous cells that comprise TNBC tumors. The issue is clear, finding a drug or a combination that is effective among two top cell lines that account for the majority of tumor cells is relatively simple, but finding one that is effective across many cell lines with different molecular phenotypes is extremely challenging. Furthermore, having a proportion of cells with a different molecular phenotype that do not respond to treatment is a good source of drug resistance development.

## Discussion

We presented evidence that supports the use of transfer learning in clinical practice for the contextualization of breast cancer patient tumor cells. We demonstrated that this procedure remains stable even when cells’ transcriptomes are characterized using a different experimental technique or when cells have a genetic mutation. We tested the efficacy of this method on a small group of breast cancer patients. The findings add to our understanding of the composition and characteristics of breast cancer tumors across multiple molecular subtypes. We predicted the most synergistic drug combinations for prioritization based on the identified compositions of the tumors. In this case, we made predictions based on patient characterizations from a small (*n* = 20) cohort. Despite the fact that single-cell RNA-seq provides unprecedented resolution, only the patient-specific descriptions and predictions should be considered facts. Population-level generalizations should be viewed as proof of concept and will almost certainly need to be reevaluated once more data from a large cohort of patients becomes available.

Drug development and testing is a lengthy process that begins with testing in cell lines that mimic the phenotypic and genetic variability of patients. We are confident that we have paved the way for data-driven drug and combination prioritization. This approach, we believe, will hasten the process of developing truly personalized (*n* = 1) pharmacological options for patients. Regardless that we used the entire dataset of cellular responses to drug combinations in this study, the authors of the dataset carefully labeled them based on the approval level for each compound. This allows for the selection of already defined as safe and approved drugs and their use in patients with limited treatment options.

Numerous unanswered questions in cancer biology could be addressed by employing a transfer learning strategy. For example, are metastatic cancer cells’ transcriptomes better represented by cell lines derived from the primary tissue or by tumors in the recipient tissue? The answer to this question will have significant clinical consequences for defining the optimal treatment strategy for patients with metastatic tumors. Given the exponential growth in the availability of single-cell data, we anticipate that this question will be addressed promptly and that this approach will be widely applied in a variety of contexts related to cancer biology.

## Data Availability

All the data and code required to replicate this study as well as the figures and tables are available at https://github.com/dosorio/scTransferLearning. The following public datasets were used in this study:

1. The transcriptionally profile of 35, 276 individual cells from 32 breast cancer cell lines reported by Gambardella, G., et al. *“A single-cell atlas of breast cancer cell lines to study tumour heterogeneity and drug response*.*”* bioRxiv (2021) (21). Accessed through: DOI, record 10.6084/m9.figshare.15022698
2. The transcriptional profile of 14, 372 wild-type cells from the MCF7 cell line reported by Ben-David, Uri, et al. *“Genetic and transcriptional evolution alters cancer cell line drug response*.*”* Nature 560.7718 (2018): 325-330 (32). Accessed through: GEO, accession number GSE114459
3. The transcriptional profile of 491 single-cells derived from the T47D cell line with a CRISPR knockout of the *CDH1* gene reported by Chen, Fangyuan, et al. *“Single-cell transcriptomic heterogeneity in invasive ductal and lobular breast cancer cells*.*”* Cancer research 81.2 (2021): 268-281 (34). Accessed through: GEO, accession number GSE144320
4. The transcriptional profile of 131 single-cells derived from the BT474 cell line and treated with 1*µ*M of Lapatinib for 10 days reported by Oren, Yaara, et al. *“Cycling cancer persister cells arise from lineages with distinct programs*.*”* Nature 596.7873 (2021): 576-582 (33). Accessed through: GEO, accession number GSE150949
5. The transcriptional profile of 130, 246 annotated single cells from 26 primary tumors reported by Wu, Sunny Z., et al. *“A single-cell and spatially resolved atlas of human breast cancers*.*”* Nature genetics 53.9 (2021): 1334-1347 (24). Accessed through: GEO, accession number GSE176078
6. The synergistic effect, potency and efficacy of 2, 025 clinically relevant two-drug combinations for 51 breast cancer, 45 colorectal and 29 pancreatic cell lines reported by Jaaks, Patricia, et al. *“Effective drug combinations in breast, colon and pancreatic cancer cells*.*”* Nature (2022): 1-8 (28). Accessed through: DOI, record 10.6084/m9.figshare.19141928

## ACKNOWLEDGEMENTS

This work was supported by the Biomedical Research Computing Facility of the University of Texas at Austin.

## AUTHOR CONTRIBUTIONS

Conceptualization: DO and SSY. Methodology: DO, DJM, and SSY. Validation: DO. Resources: DO and SSY. Data Curation: DO. Writing - Original Draft: DO. Writing - Review & Editing: DO, DJM, NS, and SSY. Supervision: SSY. Project administration: SSY. Funding acquisition: SSY, SN.

## Supplementary Material

## Supplementary Figures

**Fig. S1.**
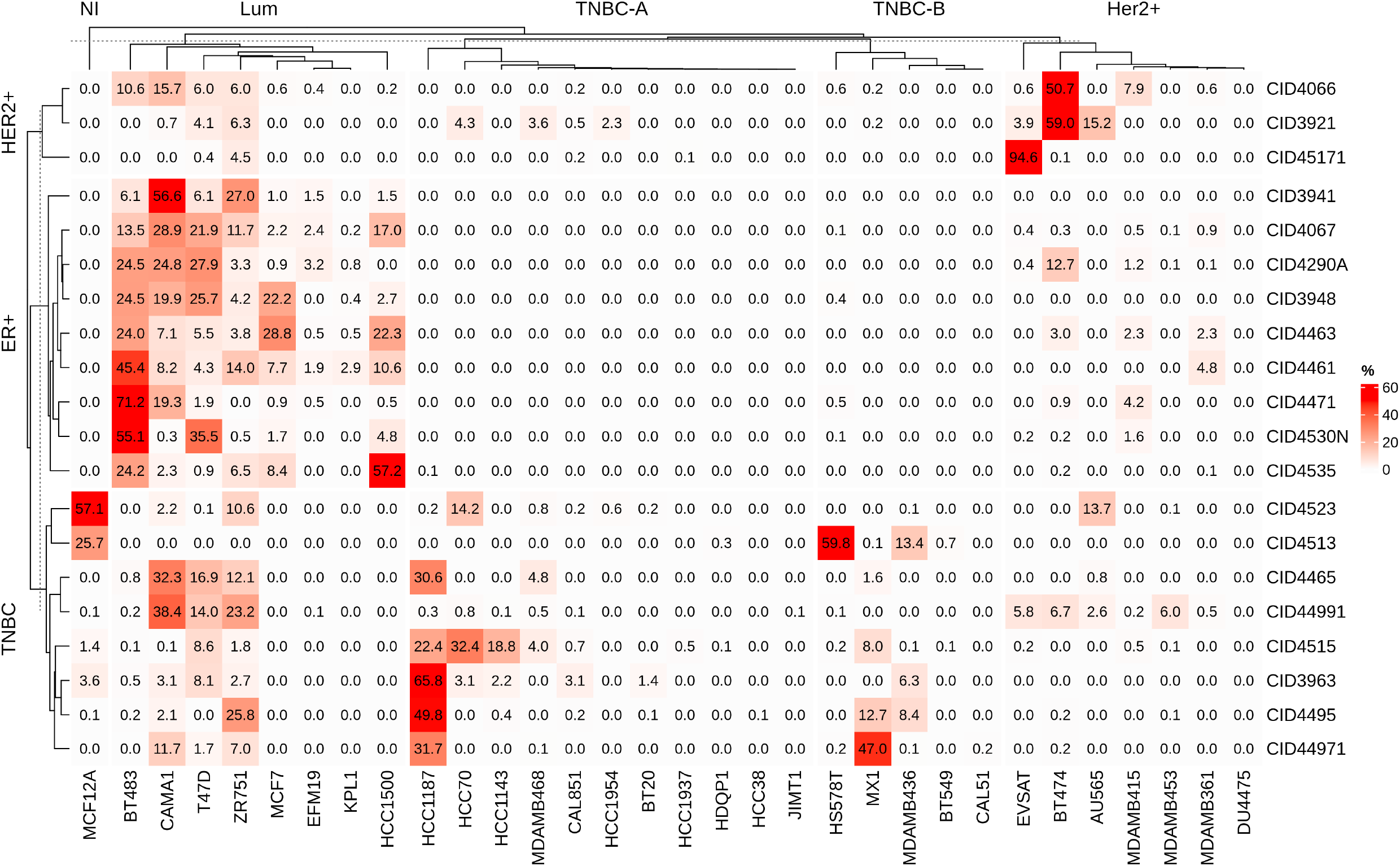
Determination of patient’s tumor composition. Composition of malignant tumor cells, expressed as a proportion (rounded to two decimals) of cell lines, for twenty patients described in the breast cancer cell atlas.

**Fig. S2.**
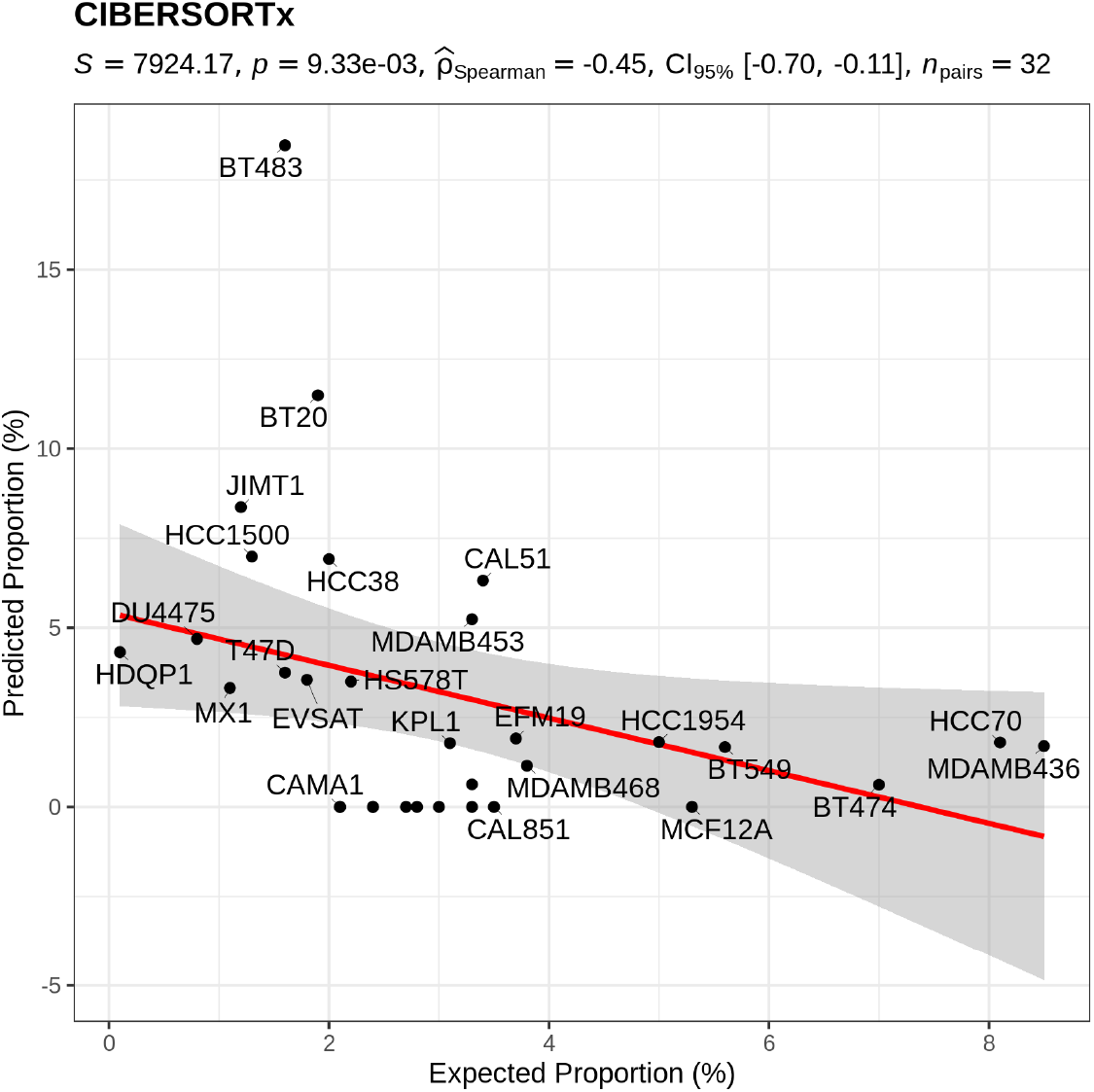
Evaluation of the precision of CIBERSORTx’s deconvolution proportions

**Fig. S3.**
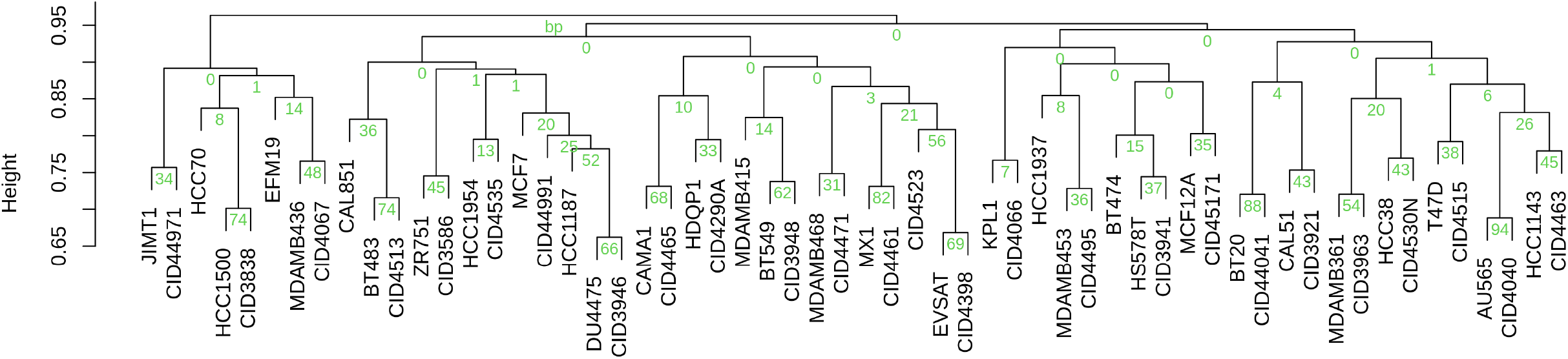
Hierarchical clustering of computed pseudo-bulk profiles using all cells. Patient profiles (CID*) are generated using all cells reported in the atlas. The percentage of bootstraps (bp) in which the relationship is recovered is indicated in green.

**Fig. S4.**
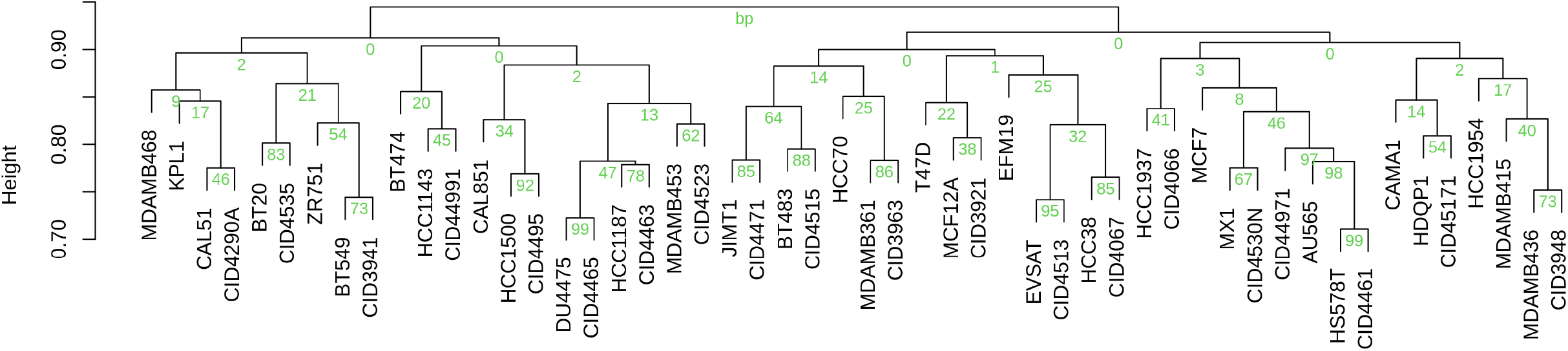
Hierarchical clustering of computed pseudo-bulk profiles using only cancer cells. Patient profiles (CID*) are generated using only the cancer cells reported in the atlas. The percentage of bootstraps (bp) in which the relationship is recovered is indicated in green.

**Fig. S5.**
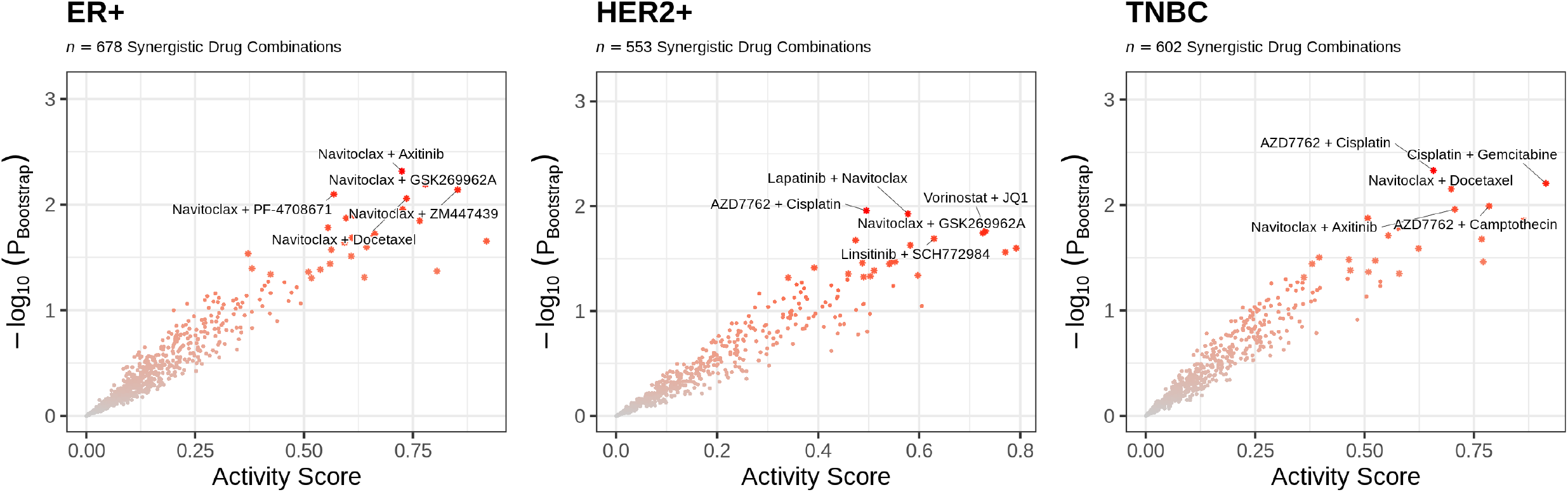
Prioritization of drug combinations exhibiting synergistic effects using uniform proportions of cell lines. **(A)** Computed for ER-positive cell lines. **(B)** Computed for HER2-positive cell lines. **(C)** Computed for TNBC cell lines.

**Fig. S6.**
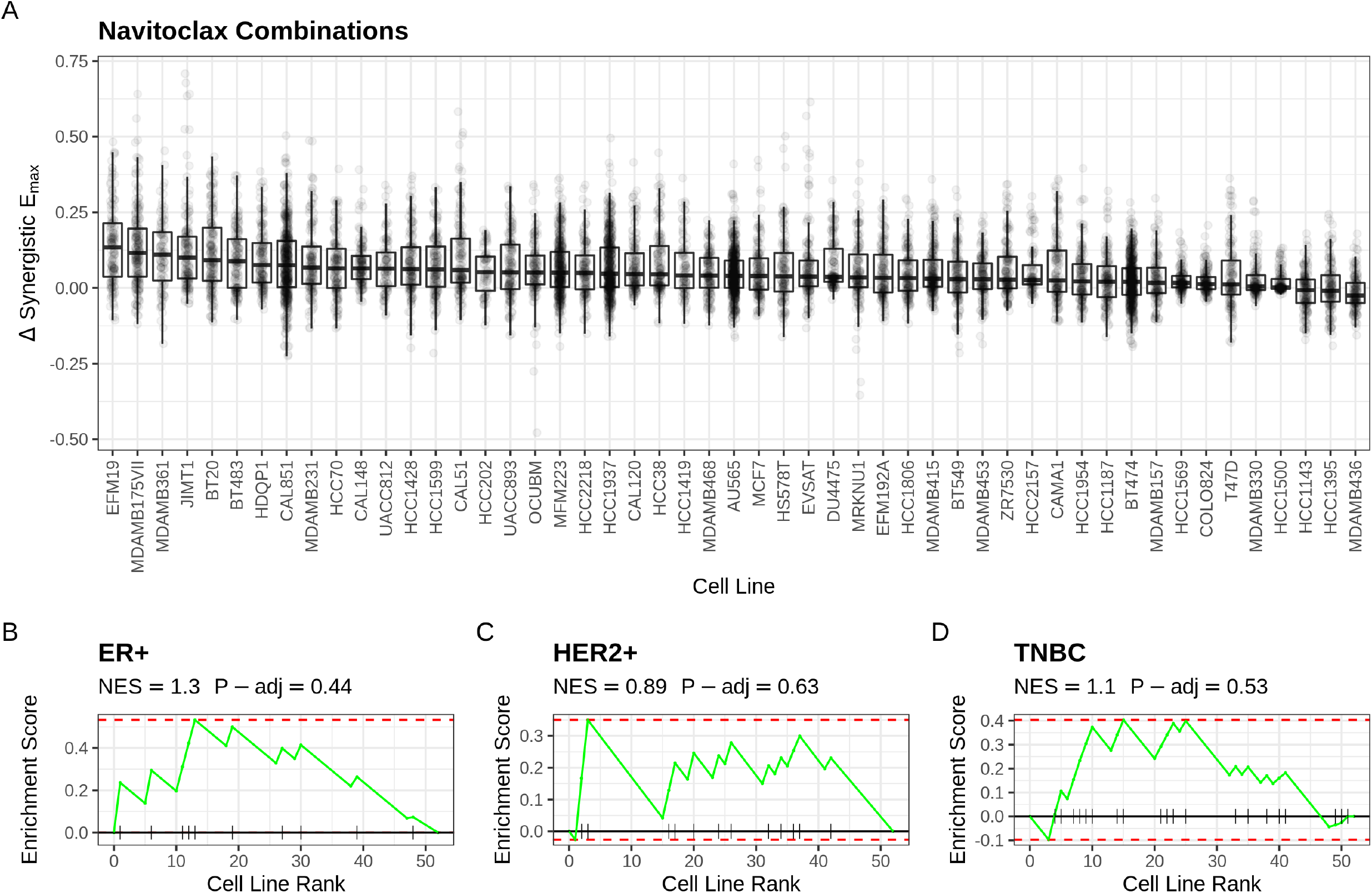
Cellular responses to Navoticlax combinations. **(A)** Distribution of the synergistic effect of Navoticlax combinations across breast cancer cell lines. **(B)** Enrichment score for ER^+^ cell lines, **(C)** Enrichment score for HER2^+^ cell lines, **(D)** Enrichment score for TNBC cell lines.

## Bibliography

1. Eoghan R Malone, Marc Oliva, Peter JB Sabatini, Tracy L Stockley, and Lillian L Siu. Molecular profiling for precision cancer therapies. Genome medicine, 12(1):1–19, 2020.

2. Jennifer I Hare, Twan Lammers, Marianne B Ashford, Sanyogitta Puri, Gert Storm, and Simon T Barry. Challenges and strategies in anti-cancer nanomedicine development: An industry perspective. Advanced drug delivery reviews, 108:25–38, 2017.

3. Girieca Lorusso and Curzio Rüegg. The tumor microenvironment and its contribution to tumor evolution toward metastasis. Histochemistry and cell biology, 130(6):1091–1103, 2008.

4. Roman Rouzier, Charles M Perou, W Fraser Symmans, Nuhad Ibrahim, Massimo Cristofanilli, Keith Anderson, Kenneth R Hess, James Stec, Mark Ayers, Peter Wagner, et al. Breast cancer molecular subtypes respond differently to preoperative chemotherapy. Clinical cancer research, 11(16):5678–5685, 2005.

5. Aleix Prat, Estela Pineda, Barbara Adamo, Patricia Galván, Aranzazu Fernández, Lydia Gaba, Marc Díez, Margarita Viladot, Ana Arance, and Montserrat Muñoz. Clinical implications of the intrinsic molecular subtypes of breast cancer. The Breast, 24:S26–S35, 2015.

6. Caicun Zhou, Yi-Long Wu, Gongyan Chen, Jifeng Feng, Xiao-Qing Liu, Changli Wang, Shucai Zhang, Jie Wang, Songwen Zhou, Shengxiang Ren, et al. Erlotinib versus chemotherapy as first-line treatment for patients with advanced EGFR mutation-positive non-small-cell lung cancer (OPTIMAL, CTONG-0802): a multicentre, open-label, randomised, phase 3 study. The lancet oncology, 12(8):735–742, 2011.

7. Jason K Sicklick, Shumei Kato, Ryosuke Okamura, Maria Schwaederle, Michael E Hahn, Casey B Williams, Pradip De, Amy Krie, David E Piccioni, Vincent A Miller, et al. Molecular profiling of cancer patients enables personalized combination therapy: the I-PREDICT study. Nature medicine, 25(5):744–750, 2019.

8. Keith T Flaherty, Robert J Gray, Alice P Chen, Shuli Li, Lisa M McShane, David Patton, Stanley R Hamilton, P Mickey Williams, A John Iafrate, Jeffrey Sklar, et al. Molecular landscape and actionable alterations in a genomically guided cancer clinical trial: National Cancer Institute Molecular Analysis for Therapy Choice (NCI-MATCH). American Society of Clinical Oncology, 2020.

9. Sameek Roychowdhury, Matthew K Iyer, Dan R Robinson, Robert J Lonigro, Yi-Mi Wu, Xuhong Cao, Shanker Kalyana-Sundaram, Lee Sam, O Alejandro Balbin, Michael J Quist, et al. Personalized oncology through integrative high-throughput sequencing: a pilot study. Science translational medicine, 3(111):111ra121–111ra121, 2011.

10. Jordi Rodon, Jean-Charles Soria, Raanan Berger, Wilson H Miller, Eitan Rubin, Aleksandra Kugel, Apostolia Tsimberidou, Pierre Saintigny, Aliza Ackerstein, Irene Braña, et al. Genomic and transcriptomic profiling expands precision cancer medicine: the WINTHER trial. Nature medicine, 25 (5):751–758, 2019.

11. Mariano J Alvarez, Prem S Subramaniam, Laura H Tang, Adina Grunn, Mahalaxmi Aburi, Gabrielle Rieckhof, Elena V Komissarova, Elizabeth A Hagan, Lisa Bodei, Paul A Clemons, et al. A precision oncology approach to the pharmacological targeting of mechanistic dependencies in neuroendocrine tumors. Nature genetics, 50(7):979–989, 2018.

12. Joo Sang Lee, Nishanth Ulhas Nair, Gal Dinstag, Lesley Chapman, Youngmin Chung, Kun Wang, Sanju Sinha, Hongui Cha, Dasol Kim, Alexander V Schperberg, et al. Synthetic lethality-mediated precision oncology via the tumor transcriptome. Cell, 184(9):2487–2502, 2021.

13. Gal Dinstag, Eldad D. Shulman, Efrat Elis, Doreen S. Ben-Zvi, Omer Tirosh, Eden Maimon, Isaac Meilijson, Emmanuel Elalouf, Eyal Schiff, Danh-Tai Hoang, Sanju Sinha, Nishanth Ulhas Nair, Joo Sang Lee, Alejandro A. Schäffer, Ze’ev Ronai, Dejan Juric, Andrea B. Apolo, William L. Dahut, Stanley Lipkowitz, Raanan Berger, Razelle Kurzrock, Antonios Papanicolau-Sengos, Fatima Karzai, Mark R. Gilbert, Kenneth Aldape, Padma S. Rajagopal, Tuvik Beker, Eytan Ruppin, and Ranit Aharonov. Clinically oriented prediction of patient response to targeted and immunotherapies from the tumor transcriptome. bioRxiv, 2022. doi: 10.1101/2022.02.27.481627.

14. Travers Ching, Daniel S Himmelstein, Brett K Beaulieu-Jones, Alexandr A Kalinin, Brian T Do, Gregory P Way, Enrico Ferrero, Paul-Michael Agapow, Michael Zietz, Michael M Hoffman, et al. Opportunities and obstacles for deep learning in biology and medicine. Journal of The Royal Society Interface, 15(141):20170387, 2018.

15. Lacey E Dobrolecki, Susie D Airhart, Denis G Alferez, Samuel Aparicio, Fariba Behbod, Mohamed Bentires-Alj, Cathrin Brisken, Carol J Bult, Shirong Cai, Robert B Clarke, et al. Patient-derived xenograft (pdx) models in basic and translational breast cancer research. Cancer and Metastasis Reviews, 35(4):547–573, 2016.

16. Peppino Mirabelli, Luigi Coppola, and Marco Salvatore. Cancer cell lines are useful model systems for medical research. Cancers, 11(8):1098, 2019.

17. Marcelo L Larramendy, Tamara Lushnikova, Anna-Maria Björkqvist, Ignacio I Wistuba, Arvind K Virmani, Narayan Shivapurkar, Adi F Gazdar, and Sakari Knuutila. Comparative genomic hybridization reveals complex genetic changes in primary breast cancer tumors and their cell lines. Cancer genetics and cytogenetics, 119(2):132–138, 2000.

18. Nifang Niu and Liewei Wang. In vitro human cell line models to predict clinical response to anti-cancer drugs. Pharmacogenomics, 16(3):273–285, 2015.

19. Nicholas E Navin. The first five years of single-cell cancer genomics and beyond. Genome research, 25(10):1499–1507, 2015.

20. Rapolas Zilionis, Camilla Engblom, Christina Pfirschke, Virginia Savova, David Zemmour, Hatice D Saatcioglu, Indira Krishnan, Giorgia Maroni, Claire V Meyerovitz, Clara M Kerwin, et al. Single-cell transcriptomics of human and mouse lung cancers reveals conserved myeloid populations across individuals and species. Immunity, 50(5):1317–1334, 2019.

21. G Gambardella, G Viscido, B Tumaini, A Isacchi, R Bosotti, and D di Bernardo. A single-cell atlas of breast cancer cell lines to study tumour heterogeneity and drug response. Nature Communications, 13(1714), 2022. doi: 10.1038/s41467-022-29358-6.

22. Evan Z Macosko, Anindita Basu, Rahul Satija, James Nemesh, Karthik Shekhar, Melissa Goldman, Itay Tirosh, Allison R Bialas, Nolan Kamitaki, Emily M Martersteck, et al. Highly parallel genome-wide expression profiling of individual cells using nanoliter droplets. Cell, 161(5):1202–1214, 2015.

23. Grace XY Zheng, Jessica M Terry, Phillip Belgrader, Paul Ryvkin, Zachary W Bent, Ryan Wilson, Solongo B Ziraldo, Tobias D Wheeler, Geoff P McDermott, Junjie Zhu, et al. Massively parallel digital transcriptional profiling of single cells. Nature communications, 8(1):1–12, 2017.

24. Sunny Z Wu, Ghamdan Al-Eryani, Daniel Lee Roden, Simon Junankar, Kate Harvey, Alma Andersson, Aatish Thennavan, Chenfei Wang, James R Torpy, Nenad Bartonicek, et al. A single-cell and spatially resolved atlas of human breast cancers. Nature genetics, 53(9):1334–1347, 2021.

25. Sinno Jialin Pan and Qiang Yang. A survey on transfer learning. IEEE Transactions on knowledge and data engineering, 22(10):1345–1359, 2009.

26. Joyce B Kang, Aparna Nathan, Kathryn Weinand, Fan Zhang, Nghia Millard, Laurie Rumker, D Moody, Ilya Korsunsky, and Soumya Raychaudhuri. Efficient and precise single-cell reference atlas mapping with Symphony. Nature communications, 12(1):1–21, 2021.

27. Mohammad Lotfollahi, Mohsen Naghipourfar, Malte D Luecken, Matin Khajavi, Maren Büttner, Marco Wagenstetter, Žiga Avsec, Adam Gayoso, Nir Yosef, Marta Interlandi, et al. Mapping single-cell data to reference atlases by transfer learning. Nature Biotechnology, 40(1):121–130, 2022.

28. Patricia Jaaks, Elizabeth A Coker, Daniel J Vis, Olivia Edwards, Emma F Carpenter, Simonetta M Leto, Lisa Dwane, Francesco Sassi, Howard Lightfoot, Syd Barthorpe, et al. Effective drug combinations in breast, colon and pancreatic cancer cells. Nature, pages 1–8, 2022.

29. Kevin L Howe, Premanand Achuthan, James Allen, Jamie Allen, Jorge Alvarez-Jarreta, M Ridwan Amode, Irina M Armean, Andrey G Azov, Ruth Bennett, Jyothish Bhai, et al. Ensembl 2021. Nucleic acids research, 49(D1):D884–D891, 2021.

30. Ilya Korsunsky, Nghia Millard, Jean Fan, Kamil Slowikowski, Fan Zhang, Kevin Wei, Yuriy Baglaenko, Michael Brenner, Po-ru Loh, and Soumya Raychaudhuri. Fast, sensitive and accurate integration of single-cell data with harmony. Nature methods, 16(12):1289–1296, 2019.

31. Max Kuhn. Building predictive models in r using the caret package. Journal of statistical software, 28:1–26, 2008.

32. Uri Ben-David, Benjamin Siranosian, Gavin Ha, Helen Tang, Yaara Oren, Kunihiko Hinohara, Craig A Strathdee, Joshua Dempster, Nicholas J Lyons, Robert Burns, et al. Genetic and transcriptional evolution alters cancer cell line drug response. Nature, 560(7718):325–330, 2018.

33. Yaara Oren, Michael Tsabar, Michael S Cuoco, Liat Amir-Zilberstein, Heidie F Cabanos, Jan-Christian Hütter, Bomiao Hu, Pratiksha I Thakore, Marcin Tabaka, Charles P Fulco, et al. Cycling cancer persister cells arise from lineages with distinct programs. Nature, 596(7873):576–582, 2021.

34. Fangyuan Chen, Kai Ding, Nolan Priedigkeit, Ashuvinee Elangovan, Kevin M Levine, Neil Carleton, Laura Savariau, Jennifer M Atkinson, Steffi Oesterreich, and Adrian V Lee. Single-cell transcriptomic heterogeneity in invasive ductal and lobular breast cancer cells. Cancer research, 81(2):268–281, 2021.

35. Gennaro Gambardella and Diego Di Bernardo. A tool for visualization and analysis of single-cell rna-seq data based on text mining. Frontiers in genetics, page 734, 2019.

36. Linda W Engel, Nathaniel A Young, Tommie Sue Tralka, Marc E Lippman, Stephen J O’Brien, and Mary Jo Joyce. Establishment and characterization of three new continuous cell lines derived from human breast carcinomas. Cancer research, 38(10):3352–3364, 1978.

37. J Kurebayashi, M Kurosumi, and H Sonoo. A new human breast cancer cell line, KPL-1 secretes tumour-associated antigens and grows rapidly in female athymic nude mice. British journal of cancer, 71(4):845–853, 1995.

38. Amanda Capes-Davis, George Theodosopoulos, Isobel Atkin, Hans G Drexler, Arihiro Kohara, Roderick AF MacLeod, John R Masters, Yukio Nakamura, Yvonne A Reid, Roger R Reddel, et al. Check your cultures! a list of cross-contaminated or misidentified cell lines. International journal of cancer, 127(1):1–8, 2010.

39. Mark Yarchoan, Lee A Albacker, Alexander C Hopkins, Meagan Montesion, Karthikeyan Murugesan, Teena T Vithayathil, Neeha Zaidi, Nilofer S Azad, Daniel A Laheru, Garrett M Frampton, et al. Pd-l1 expression and tumor mutational burden are independent biomarkers in most cancers. JCI insight, 4(6), 2019.

40. Francisco Beca and Kornelia Polyak. Intratumor heterogeneity in breast cancer. Novel biomarkers in the continuum of breast cancer, pages 169–189, 2016.

41. Jørgen Fogh, William C Wright, and James D Loveless. Absence of hela cell contamination in 169 cell lines derived from human tumors. Journal of the National Cancer Institute, 58(2):209–214, 1977.

42. Etienne Y Lasfargues, William G Coutinho, and Ernest S Redfield. Isolation of two human tumor epithelial cell lines from solid breast carcinomas. Journal of the National Cancer Institute, 61(4): 967–978, 1978.

43. Alexandre F Aissa, Abul BMMK Islam, Majd M Ariss, Cammille C Go, Alexandra E Rader, Ryan D Conrardy, Alexa M Gajda, Carlota Rubio-Perez, Klara Valyi-Nagy, Mary Pasquinelli, et al. Single-cell transcriptional changes associated with drug tolerance and response to combination therapies in cancer. Nature communications, 12(1):1–25, 2021.

44. Ibiayi Dagogo-Jack and Alice T Shaw. Tumour heterogeneity and resistance to cancer therapies. Nature reviews Clinical oncology, 15(2):81–94, 2018.

45. Philippe L Bedard, Aaron R Hansen, Mark J Ratain, and Lillian L Siu. Tumour heterogeneity in the clinic. Nature, 501(7467):355–364, 2013.

46. Ana C Garrido-Castro, Nancy U Lin, and Kornelia Polyak. Insights into molecular classifications of triple-negative breast cancer: improving patient selection for treatment. Cancer discovery, 9(2): 176–198, 2019.

47. Xiaofeng Dai, Hongye Cheng, Zhonghu Bai, and Jia Li. Breast cancer cell line classification and its relevance with breast tumor subtyping. Journal of Cancer, 8(16):3131, 2017.

48. Jessica Vamathevan, Dominic Clark, Paul Czodrowski, Ian Dunham, Edgardo Ferran, George Lee, Bin Li, Anant Madabhushi, Parantu Shah, Michaela Spitzer, et al. Applications of machine learning in drug discovery and development. Nature reviews Drug discovery, 18(6):463–477, 2019.

49. Hannah K Weir, Robert N Anderson, Sallyann M Coleman King, Ashwini Soman, Trevor D Thompson, Yuling Hong, Bjorn Moller, and Steven Leadbetter. Heart disease and cancer deaths—trends and projections in the united states, 1969–2020. Preventing chronic disease, 13, 2016.

50. DCFR Koboldt, Robert Fulton, Michael McLellan, Heather Schmidt, Joelle Kalicki-Veizer, Joshua McMichael, Lucinda Fulton, David Dooling, Li Ding, Elaine Mardis, et al. Comprehensive molecular portraits of human breast tumours. Nature, 490(7418):61–70, 2012.

51. Chloé B Steen, Chih Long Liu, Ash A Alizadeh, and Aaron M Newman. Profiling cell type abundance and expression in bulk tissues with cibersortx. In Stem Cell Transcriptional Networks, pages 135–157. Springer, 2020.

52. Brandon Jew, Marcus Alvarez, Elior Rahmani, Zong Miao, Arthur Ko, Kristina M Garske, Jae Hoon Sul, Kirsi H Pietiläinen, Päivi Pajukanta, and Eran Halperin. Accurate estimation of cell composition in bulk expression through robust integration of single-cell information. Nature communications, 11(1):1–11, 2020.

53. Charles Spearman. The proof and measurement of association between two things. 1961.

54. Ryota Suzuki and Hidetoshi Shimodaira. Pvclust: an r package for assessing the uncertainty in hierarchical clustering. Bioinformatics, 22(12):1540–1542, 2006.

55. Thomas J Gonda and Robert G Ramsay. Directly targeting transcriptional dysregulation in cancer. Nature Reviews Cancer, 15(11):686–694, 2015.

56. N. Chatterjee and T. G. Bivona. Polytherapy and targeted cancer drug resistance. Trends Cancer, 5(3):170–182, 2019. ISSN 2405-8025 (Electronic) 2405-8025 (Linking). doi: 10.1016/j.trecan.2019.02.003.

57. J Zhang, Y Lin, XJ Sun, BY Wang, ZH Wang, JF Luo, LP Wang, S Zhang, J Cao, ZH Tao, et al. Biomarker assessment of the cbcsg006 trial: a randomized phase iii trial of cisplatin plus gemcitabine compared with paclitaxel plus gemcitabine as first-line therapy for patients with metastatic triple-negative breast cancer. Annals of Oncology, 29(8):1741–1747, 2018.

58. Christin Tse, Alexander R Shoemaker, Jessica Adickes, Mark G Anderson, Jun Chen, Sha Jin, Eric F Johnson, Kennan C Marsh, Michael J Mitten, Paul Nimmer, et al. Abt-263: a potent and orally bioavailable bcl-2 family inhibitor. Cancer research, 68(9):3421–3428, 2008.

59. Dong-Joon Min, Siping He, and Jeffrey E Green. Birinapant (tl32711) improves responses to gem/azd7762 combination therapy in triple-negative breast cancer cell lines. Anticancer research, 36(6):2649–2657, 2016.

60. Edward Sausville, Patricia LoRusso, Michael Carducci, Judith Carter, Mary F Quinn, Lisa Malburg, Nilofer Azad, David Cosgrove, Richard Knight, Peter Barker, et al. Phase i dose-escalation study of azd7762, a checkpoint kinase inhibitor, in combination with gemcitabine in us patients with advanced solid tumors. Cancer chemotherapy and pharmacology, 73(3):539–549, 2014.

61. S Nylander, B Kull, J. Björkman, JC Ulvinge, N Oakes, BM Emanuelsson, M Andersson, T Skärby, T Inghardt, O Fjellström, et al. Human target validation of phosphoinositide 3-kinase (pi3k) β: effects on platelets and insulin sensitivity, using azd6482 a novel pi3kβ inhibitor. Journal of Thrombosis and Haemostasis, 10(10):2127–2136, 2012.

62. Junjun Zhang, Joachim Baran, Anthony Cros, Jonathan M Guberman, Syed Haider, Jack Hsu, Yong Liang, Elena Rivkin, Jianxin Wang, Brett Whitty, et al. International cancer genome consortium data portal—a one-stop shop for cancer genomics data. Database, 2011, 2011.

63. Miriam Martini, Maria Chiara De Santis, Laura Braccini, Federico Gulluni, and Emilio Hirsch. Pi3k/akt signaling pathway and cancer: an updated review. Annals of medicine, 46(6):372–383, 2014.

